# Open Imputation Server provides secure Imputation services with provable genomic privacy

**DOI:** 10.1101/2021.09.30.462262

**Authors:** Arif O. Harmanci, Miran Kim, Su Wang, Wentao Li, Yongsoo Song, Kristin E. Lauter, Xiaoqian Jiang

**Affiliations:** Center for Precision Health, School of Biomedical Informatics, University of Texas Health Science Center, Houston, TX, 77030, USA; Department of Computer Science and Engineering and Graduate School of Artificial Intelligence, Ulsan National Institute of Science and Technology, Ulsan, 44919, Republic of Korea; Center for Secure Artificial intelligence For hEalthcare (SAFE), School of Biomedical Informatics, University of Texas Health Science Center, Houston, TX, 77030, USA; Department of Computer Science and Engineering, Seoul National University, Seoul, 08826, Republic of Korea; West Coast Head of Research Science, Facebook AI Research (FAIR), Seattle, WA, USA

## Abstract

**Summary:** As DNA sequencing data is available for personal use, genomic privacy is becoming a major challenge. Nevertheless, high-throughput genomic data analysis outsourcing is performed using pipelines that tend to overlook these challenges.

**Results:** We present a client-server-based outsourcing framework for genotype imputation, an important step in genomic data analyses. Genotype data is encrypted by the client and encrypted data are used by the server that never observes the data in plain. Cloud-based framework can benefit from virtually unlimited computational resources while providing provable confidentiality. We demonstrate server’s utility from several aspects using genotype dataset from the 1000 Genomes datasets. First, we benchmark the accuracy of common variant imputation in comparison to BEAGLE, a state-of-the-art imputation method. We also provide the detailed time requirements of the server to showcase scaling of time usage in different steps of imputation. We also present a simple correlation metric that can be used to estimate imputation accuracy using only the reference panels. This is important for filtering the variants in downstream analyses. As a further demonstration and a different use case, we performed a simulated genomewide association study (GWAS) using imputed and known genotypes and highlight potential utility of the server for association studies. Overall, our study present multiple lines of evidence for usability of secure imputation service.

**Availability:** Server is publicly available at https://www.secureomics.org/OpenImpute. Users can anonymously test and use imputation server without registration.

## Background

DNA sequencing has entered many aspects of life including recreational purposes, clinical use, and research[1]. Numerous compute-heavy genomic analysis tasks are outsourced to servers. Untrusted servers may make it challenging to provide confidentiality of the data as a result of unauthorized access (e.g. hacking) [2]. While secure data analysis methods are available, bioinformatics is slow to adopt these mainly due to concerns about practicality. Here, we focus on genotype imputation[3], a computationally intensive task where a sparse set of genotyped variants (tag variants) are used to impute genotypes of untyped variants that are in a reference genotype panel such as The 1000 Genomes Project[4]. These methods have shown great promise for decreasing cost in population-scale association studies whereby low-cost genotyping arrays are used to genotype tag variants, which is followed by in-silico imputation of remaining sets of variants. Numerous servers are established for outsourcing the genotype imputation[5]. Here, we present a secure client-server framework that provides confidentiality of the genotype data by the homomorphic encryption (HE) schemes[6] that provably guarantee the confidentiality of data. Genotype data is encrypted once at the client and submitted to the server, which securely imputes the untyped variants without decrypting the genotypes. The imputation is performed by secure evaluation of machine learning models that we recently developed[7]. In this manuscript, we build on top of the models and develop a simple metric that can be used to estimate genotype imputation accuracy (Allelic R2) using reference panel and imputed genotypes. To demonstrate usability, we compare accuracy of imputation and estimate how time requirements scale with increasing datasize. As a separate example, we present an application of secure imputation service for genome wide association study.

## Methods

Imputation server workflow is illustrated in Figure 1.

**Figure 1:**
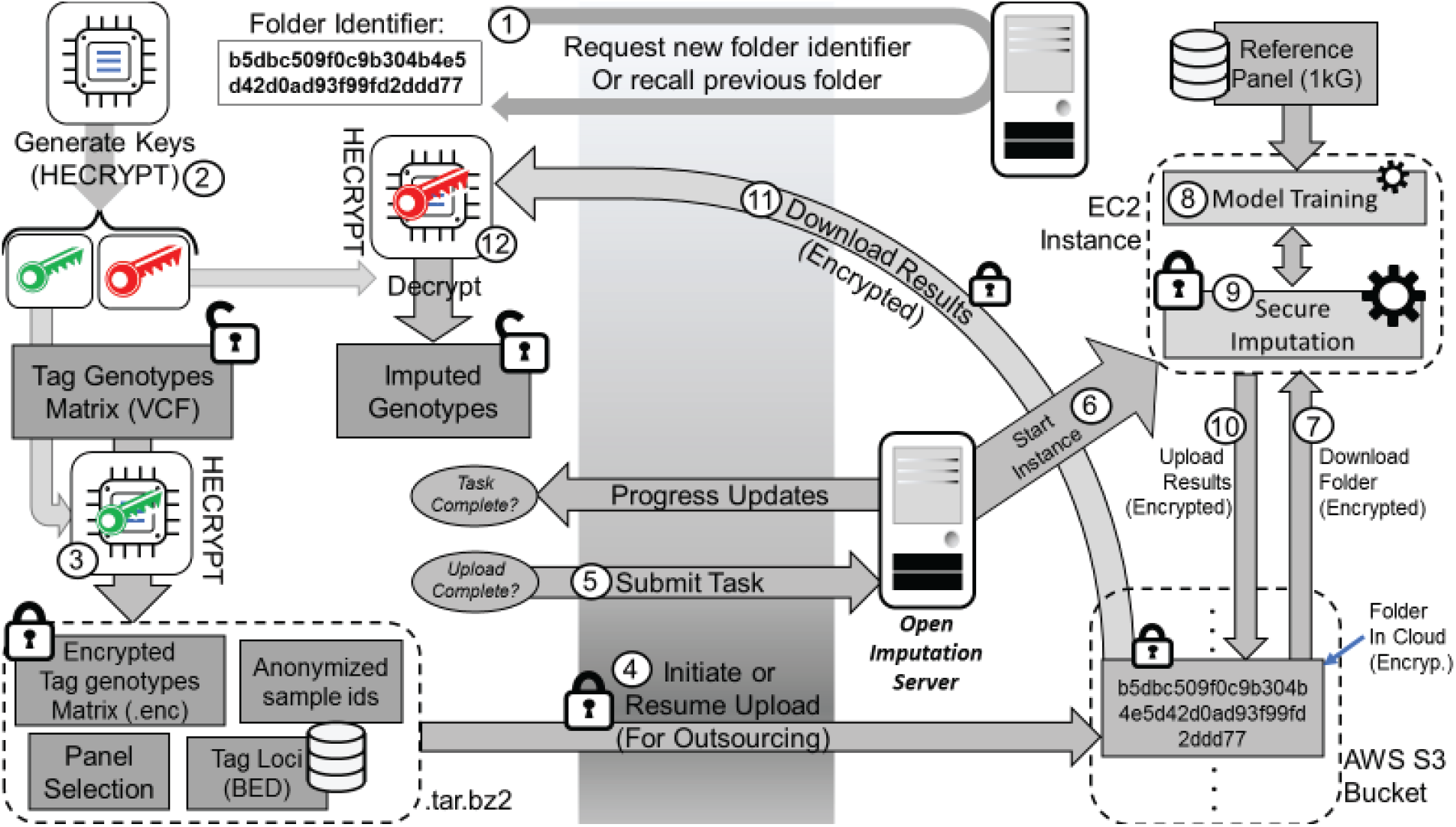
Illustration of Open Imputation Server. Imputation starts by using HECRYPT to generate public/privatekeys (shown in green and red). After the tag genotypes are encrypted, they are uploaded to S3 bucket. The server initiates an AWS compute instance, which performs model building and secure imputation. The encrypted imputation results are uploaded to the S3 bucket. The user downloads the results and decrypts them using the private key.

### Encryption of Tag Variant Genotypes

To start imputation, the client uses HECRYPT (https://github.com/harmancilab/HECRYPT) to process tag variant genotypes (Figure 1). HECRYPT is used to 1) Preprocess VCF file to check for formatting and input validation, 2) Generate public/private keys, 3) Encrypt the genotype matrix, 4) Clean the plaintext intermediate data files. The final data to be uploaded contains the encrypted tag genotype matrix, the public key, tag variant positions, and the sample size. The private key is must be kept confidential since it cannot be recovered later. The public key (“.public_key”) is necessary for encryption; it cannot be used to decrypt data. By default, HECRYPT uses 128-bit security as per Albrecht’s LWE estimator[8].

### Uploading the Encrypted Genotypes and Anonymity of Users

Encrypted data is uploaded to the server. As the encrypted genotype files tend to be quite large (roughly expansion factor of 4), the server can resume a previous upload if it is interrupted. To identify a previous upload, the server generates a folder identifier, (a key in the AWS cloud) where the client’s data is stored. This identifier has no sensitive information about the client and is uniquely generated for every new connection. The folder identifier must be recorded by the client because it cannot be recalled by anyone (even at the server-side) in case it is lost. If the folder is found by the server, its information is loaded and it can be resumed. The server keeps a simple hash of the encrypted data file to make sure the resumed file matches the original file.

The data upload is performed directly to our private AWS S3 bucket from the client (server is bypassed in data uploads) that is configured to be private with no accession from the outside world. In addition, we have disabled the usage of root user privileges (as recommended by AWS) accessible by a managed AIM account. To note, even if the bucket is fully compromised by a hacker, no privacy breaches are possible since all of the genotype information is encrypted.

In this process, there is no personal information exchange with the user. Server does not ask for any personal information (e.g., names, user ids, email addresses, passwords) that may potentially identify the user. Furthermore, the server is configured to not store or log the IP addresses of connecting clients. In case the server is not trusted, the client can use a VPN service to hide identity. The server uses TLS connections (against man-in-the-middle attacks) to the client by default.

### Submission of Imputation Task

After the data is uploaded successfully, the client submits the imputation task. The server initiates an EC2 spot instance (lower costs) to perform imputation task. The imputation instance sends task progress to the S3, which is regularly checked by the client’s browser. After imputation task is finished, a secure link is generated for the results file. The results can be downloaded using this link through the browser. HECRYPT is used to decrypt the results using the private key. An example VCF file and an encrypted genotype dataset (ready to be submitted) are available at https://github.com/harmancilab/HECRYPT with extensive documentation. Imputation task is currently performed by the default OLS-based imputation models with vicinity parameter of 32 tag variants for each untyped variant. The 1000 Genomes Panel is used for training imputation models. Only target variants with minor allele frequency above 5% are imputed. After decryption, HECRYPT can be used to estimate genotype likelihoods and generate accuracy estimates[9].

### Implementation and Compatibility

Webpage was implemented with HTML and Javascript. The server backend is implemented with PHP (AWS SDK2). The imputation pipeline is implemented as a single bash script that can be easily replaced to enable different models. Compatibility is tested with Chrome, Firefox, and SAFARI browsers.

## Results

We aim at demonstrating the utility of secure imputation server with comparisons. First, we provide accuracy benchmarks with BEAGLE algorithm[10], a state-of-the-art hidden Markov model-based genotype imputation method. We also report time usage of different steps.

### Accuracy Comparison with BEAGLE

We tested accuracy of imputation using genotype dataset from the 1000 Genomes Project[4]. We selected three different sets of tag variants (1000, 2000, 3000) chromosome 22 (Between positions 30,000,000-35,000,000) and performed imputation of common variants whose minor allele frequencies are greater than 5%. We use the variants on 1000 Genomes Project that overlap with Illumina Duo version 3 platform to present a realistic scenario. All the remaining common variants are treated as untyped target variants whose genotypes are imputed. We divided 2,504 samples randomly into 1,500 training samples and 1,004 testing samples. Training sample are used for imputation model training and testing samples are used for accuracy benchmarking. For the 3 datasets, we computed the genotype concordance (fraction of correctly imputed genotypes) and genotype R2 metric, the square of Pearson correlation between known and imputed allelic dosages. Figure 2 shows the comparison of BEAGLE. On average, the genotype concordance is slightly lower for secure imputation approach, generally between 1-2% (Fig 2a). Next we evaluated the fraction of variants that are imputed whose genotype R2 is greater than 0.5, which is indicative of reliable imputation. Figure 2b shows that both methods provide a high fraction of reliably imputed variants, although secure imputation generates slightly lower quality imputations.

**Figure 2:**
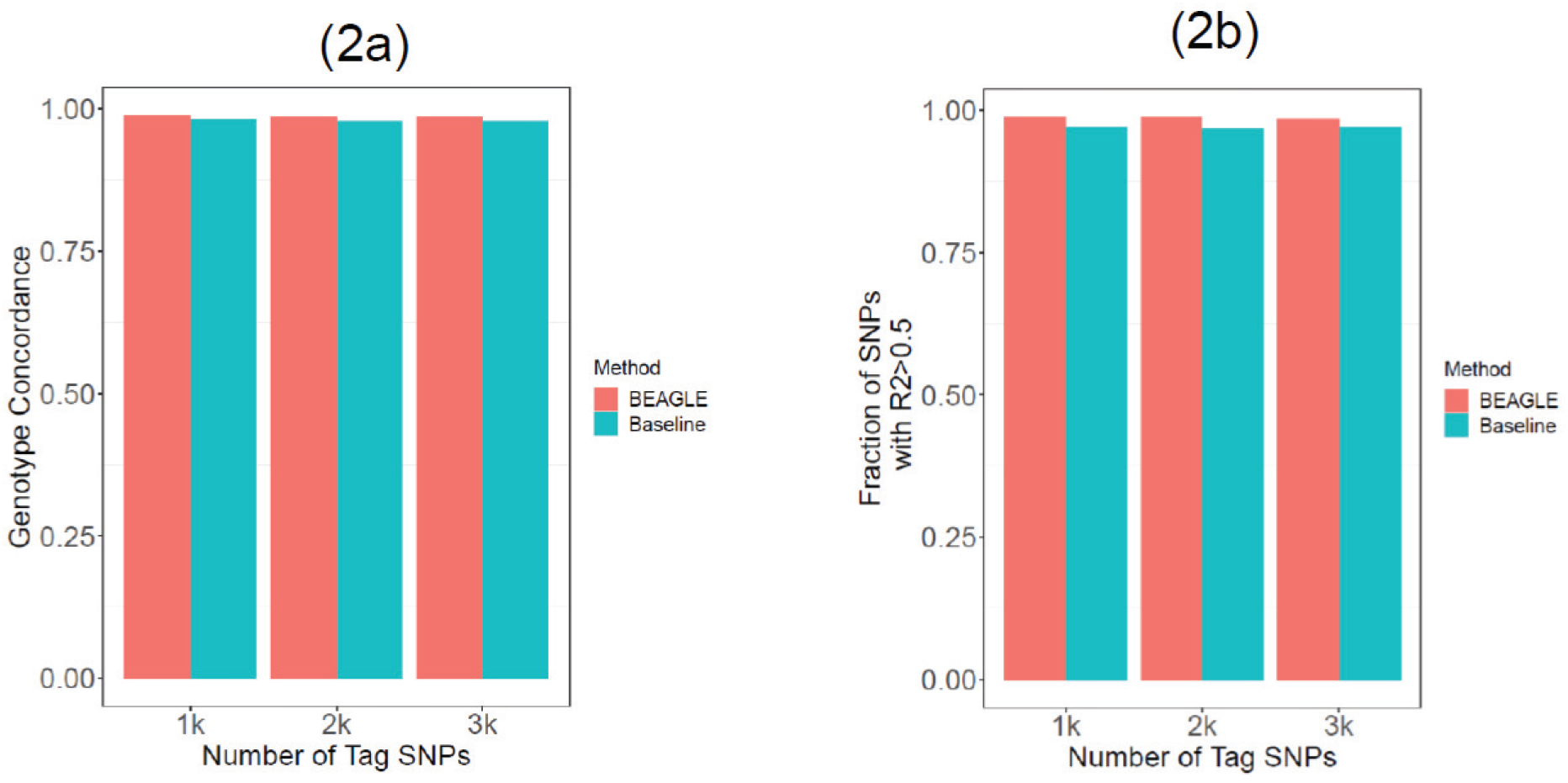
Imputation concordance (a) and fraction of reliable SNPs for 3 sets of tag SNPs (b) with BEAGLE and baseline imputation method, which is OLS models. Different tag sets are shown on the X-axis. Color indicates the method that is used for imputation.

### Estimation of Genotype R2 using Reference Panel

Genotype R2[11], square of Pearson correlation between known and imputed allelic dosages, is an important quantity to measure the imputation accuracy. We propose using the reference panel on which the imputation models are train for estimating genotype R2. This is clearly very useful since the users do not have access to true genotypes of the target variants. Having a measure of genotype R2 can be used to filter low quality imputations. We basically propose (1) Train imputation model on the reference panel, (2) Impute the target variants on the reference panel as if they are untyped, (3) Compute the square of Pearson correlation between the imputed genotypes and the known genotypes on the reference panel. We applied this metric to evaluate its effectiveness using the 3000 tag variant dataset, for which approximately 16,000 target variants are imputed. It should be noted that predicted R2 is computed on strictly different set of individuals, i.e., the predicted R2 uses training sample and true R2 is the testing individuals, whose genotypes are not known. Figure 3a shows the scatter plot for predicted genotype R2 and the true R2 metric, which is computed with known genotypes. It can be seen that there is very high concordance between the true R2 metric and the predicted R2 metric. This result indicates that our proposed metric can be used for filtering the imputed variants, which is very important for downstream analyses. One limitation of the proposed R2 metric is that it originates from reference panel dataset, i.e., the trained models are used to impute the same panel. Therefore, we expect that predicted R2 metric is an overestimate of true R2 metric. Figure 3b shows that this is indeed the case, the difference between predicted R2 and true R2 is shifted slightly towards positive values. This indicates that more research is needed to analyze our proposed R2 predictions.

**Figure 3:**
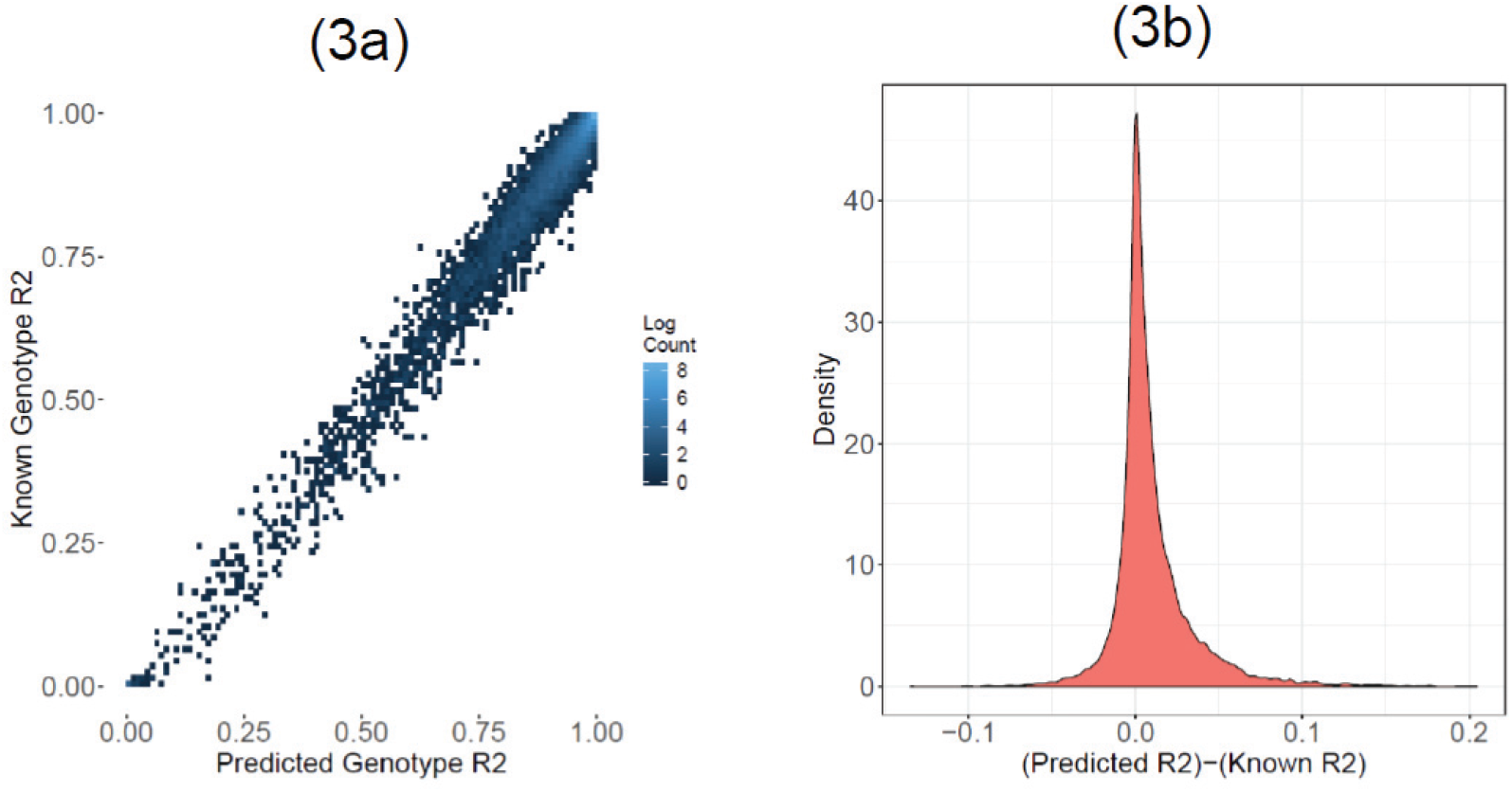
The predicted R2 comparison to true genotype R2. Scatter plot in (a) shows the distribution of predicted R2 (x-axis) vs the known R2 (y-axis). The plot in (b) shows the distribution difference between predicted R2 and known R2. Note that the distribution is skewed to right, indicating that predicted R2 overestimates true R2.

### Time Requirements of Imputation Service

An important aspect of secure imputation is the time requirements. Our methods make use of HE-based encryption scheme, which may require large compute time. To demonstrate the feasibility, we divided the run time requirements into 8 steps: (1) Preprocessing, (2) Data compression, (3) Key generation, (4) Encryption, (5) Model training, (6) Secure Evaluation, (7) Decryption, (8) Decompression. Figure 4 shows the time requirements. Other than key generation step (which take several milliseconds), all steps scale linearly with different increasing tag variant number. The largest time is required by the model building step which stays under 5 minutes. Other steps have comparable run times. Notably secure evaluation is extremely fast: For 16,000 target variants, it took 8 seconds to evaluate the trained models. This result indicates that when the models are pretrained (e.g., an existing array platform), secure imputation service can be very fast since model training step is not needed.

**Figure 4:**
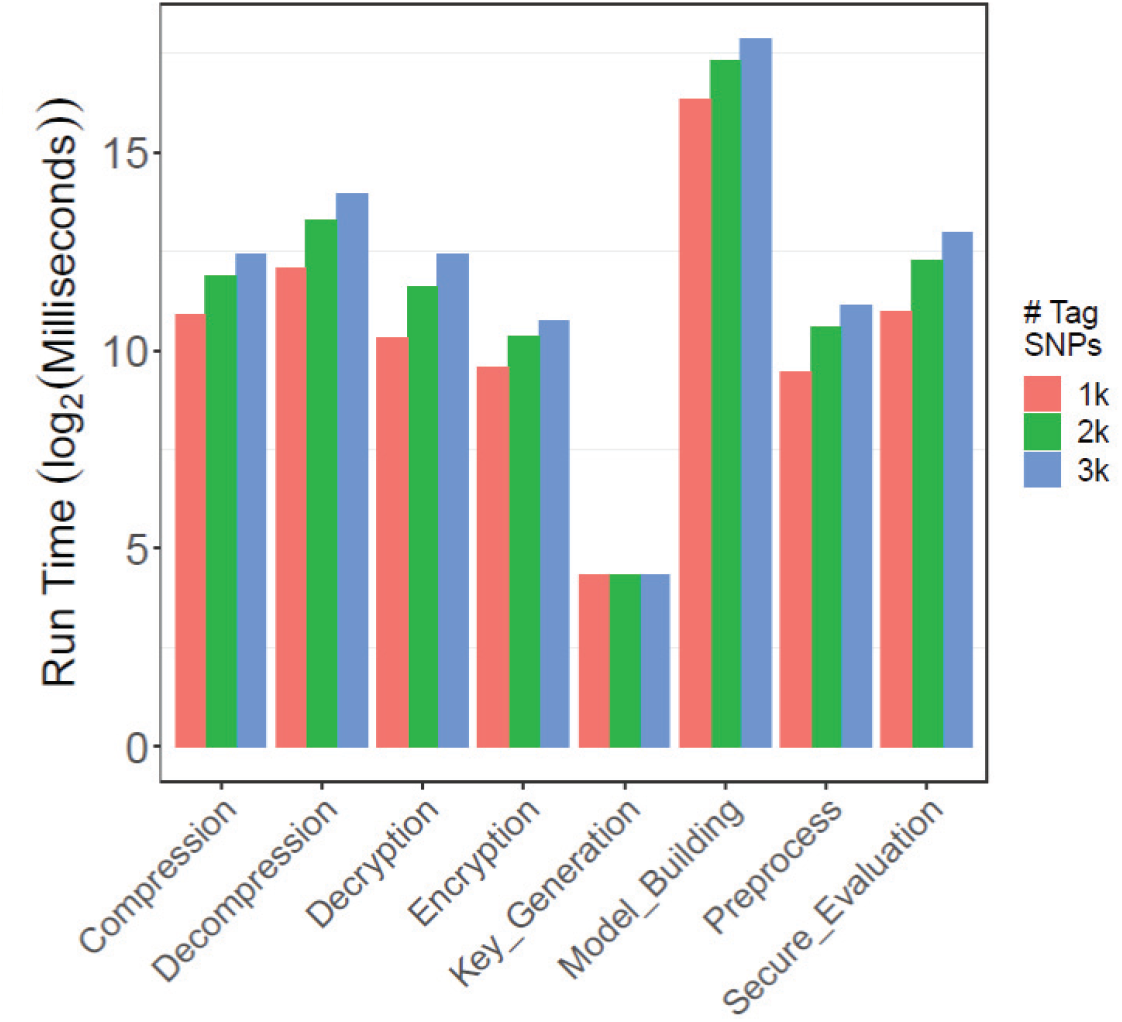
Time requirements of Imputation Server. 8 different steps are shown on the x-axis. Each tag variant set is colored differently. Y-axis shows the time requirement (logarithm of run-time in milliseconds).

### Simulated GWAS using Imputed Genotypes

As the accuracy of imputations are slightly lower than BEAGLE, we asked how much this would impact a downstream analysis. For this, we simulated a genomewide association study as a test case scenario. For this case, we focused on all target variants on chromosome 22, wherein are in total 83,072 untyped variants. We next selected 100 untyped variants as the causal variants that impact our simulated phenotype. To test the impact of allele frequency, we selected the three sets of causal variants based on allele frequency. We used variants with allele frequencies lower than 8%, 10%, and 99% (all variants) to randomly select the causal variants. For each case, we assigned a modest effect size of 0.2 to all 100 variants. To integrate population effect (not known to the GWA study), we assigned population specific effect by using higher effect size for European (EUR) and African (AFR) populations. We also assigned gender-specific effect by increasing phenotype in males. The population and gender information are extracted from the 1000 Genomes Project. After simulating phenotypes for the 3 variant sets, we used the known genotypes to identify the causal variants using plink[12]. The identified variants are our baseline GWAS results. We next performed secure imputation of the untyped variants to generate the imputed variant dataset. These variants are also input to plink to perform the testing association study. We compared the GWAS results with imputed genotypes to the baseline GWAS results.

We first compared the significance that are assigned to causal variants by baseline GWAS and GWAS with imputed genotypes. Figure 5 shows the p-value assigned to the 3 sets of causal variants. We generally observed a fairly good agreement on the assigned significance, although there are some discrepancies. As the frequency of causal variants increase, the concordance in the assigned p-values tend to increase (Figure 5a,5b,5c). We next evaluated all the 83,072 variants and sorted them with respect to the assigned increasing p-value. We compared the top scoring variants between the baseline GWAS result and GWAS with imputed genotypes based on two criteria: (1) Overlap between the SNP identifiers (2) Overlap between the vicinity of variants by looking for variants within +/-50 kilobases of the top variant. For this, we looked at first 500 variants and computed the overlap between top scoring variants using the two criteria. Figure 6 shows the top scoring variant overlap. For low frequency variants, there is generally around 50-60% overlap when matching top variants by their SNP identifiers. When we do matching by locus, we observed a much higher overlap between variants at around 80%. When the allele frequency of causal variants increase, the concordance increases as well. For locus-based matching of top candidates, we observed the overlap is frequently 100% for the top variants. These results indicate that our genotype imputation service can be beneficial for GWAS.

**Figure 5:**
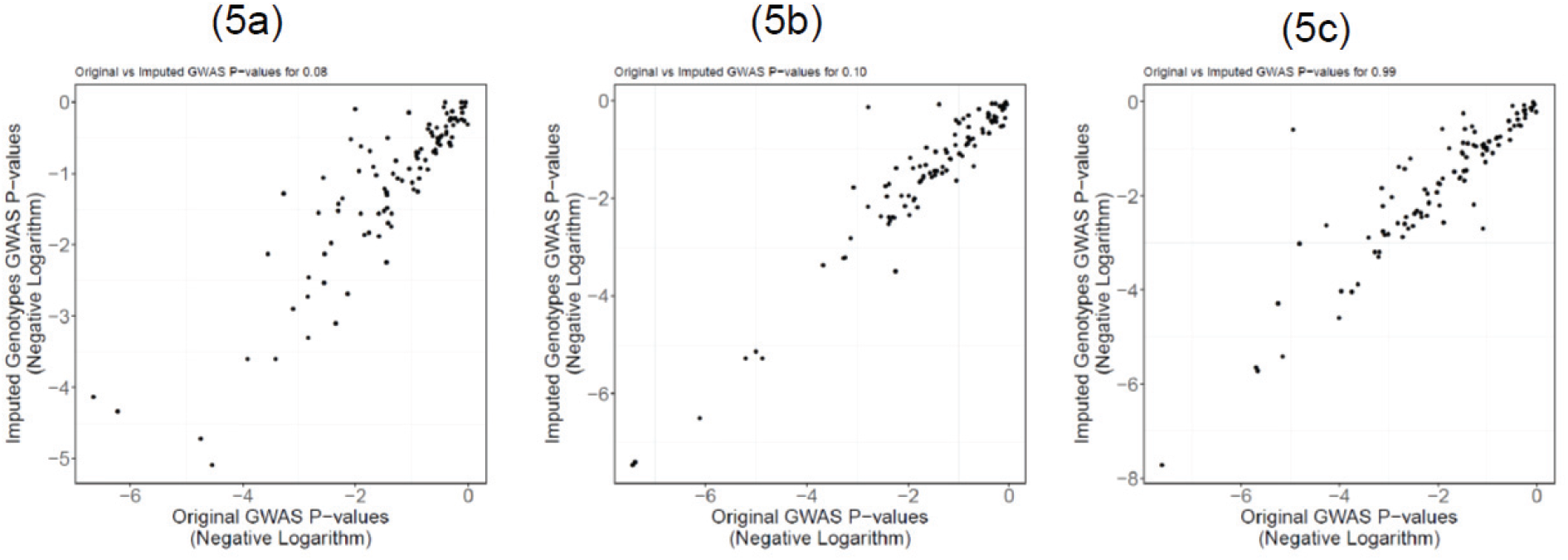
Comparison of baseline GWAS p-values (x-axis) with the GWAS p-values computed using imputed genotypes. The causal variants with allele frequency lower than 8% (a), 10% (b), and 99% (c) are shown in each plot.

**Figure 6:**
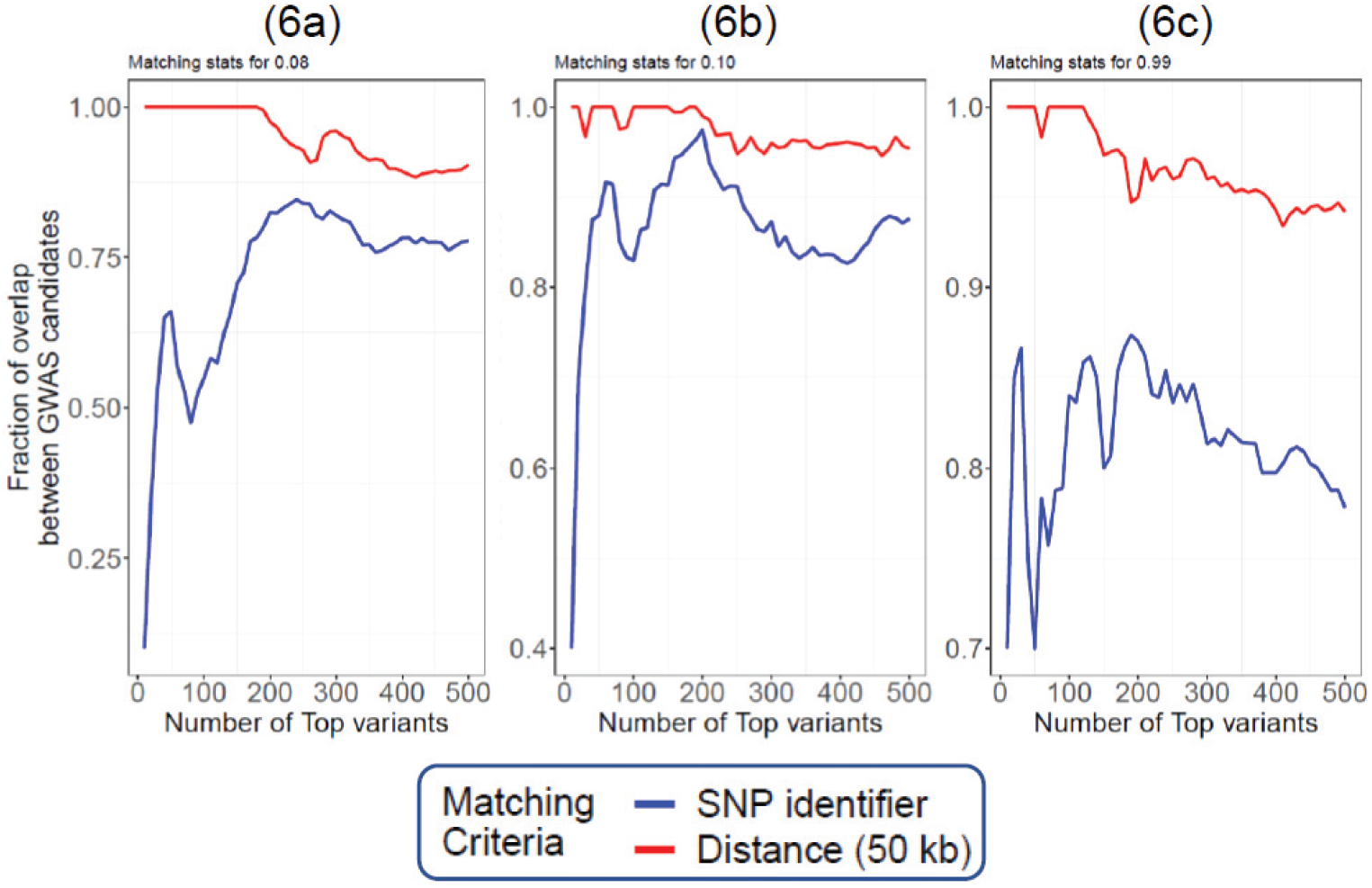
The overlap between the top candidates out of 83,072 target variants from baseline GWAS and GWAS with imputed genotypes. Each plot shows the fraction of matching variants by SNP identifier (blue) and by distance (+/-50kb vicinity) for the top variants shown in x-axis. Y-axis shows the fraction of overlap between the GWAS candidates from two approaches. The causal variants with allele frequency lower than 8% (a), 10% (b), 99% (c) are shown in the figure.

## Conclusions and Future Work

Our server demonstrates the feasibility of a client-server architecture for HE-based high-throughput genomic data analysis, which we hope can be adapted by other secure methods. We are currently extending our server to allow integration of 3^rd^ party secure genomic analysis services as docker images for demonstration purposes or high throughput applications. We are also building pre-trained imputation models that will be seamlessly integrated into imputation pipelines to make them more accurate and efficient. Our demonstration of GWAS provides potential applicability in different scenarios and downstream analyses.

## Notes

### Competing Interest Statement

The authors have declared no competing interest.

